# A polyyne toxin produced by an antagonistic bacterium blinds and lyses a green microalga

**DOI:** 10.1101/2021.03.24.436739

**Authors:** Vivien Hotter, David Zopf, Hak Joong Kim, Anja Silge, Michael Schmitt, Prasad Aiyar, Johanna Fleck, Christian Matthäus, Julian Hniopek, Qing Yan, Joyce Loper, Severin Sasso, Christian Hertweck, Jürgen Popp, Maria Mittag

## Abstract

Microalgae are key contributors to global carbon fixation and the basis of many food webs. In nature, their growth is often supported or suppressed by other microorganisms. The bacterium *Pseudomonas protegens* Pf-5 arrests the growth of the green alga *Chlamydomonas reinhardtii*, deflagellates the alga by the cyclic lipopeptide orfamide A, and alters its morphology. Using a combination of Raman microspectroscopy, genome mining and mutational analysis, we discovered a novel polyyne toxin we name protegencin that is secreted by *P. protegens* and penetrates algal cells to destroy their primitive visual system, the eyespot. Together with secreted orfamide A, protegencin prevents the phototactic behavior of *C. reinhardtii* needed to perform optimal photosynthesis. A protegencin-deficient biosynthetic mutant of *P. protegens* does not affect growth or eyespot carotenoids of *C. reinhardtii*. Thus, protegencin acts in a direct and destructive way, and reveals at least a two-pronged molecular strategy used by algicidal bacteria.

## Introduction

Photosynthetically active microorganisms, comprising cyanobacteria and eukaryotic microalgae, contribute about 50% to global CO_2_ fixation (1). As primary producers, they are fundamental to food webs (2, 3). Algal activities can also influence biogeochemical processes as exemplified recently with the Greenland ice sheet (4). In nature, microalgae are usually associated with other microbes that influence their fitness in either mutualistic or antagonistic interactions (3, 5, 6) and the exchange of natural products can play a central role (7). Despite their ecological importance, the interactions of microalgae with other microorganisms are still poorly understood at the molecular level, especially relative to our understanding of higher plant-microbe interactions (8).

In recent years, the unicellular, biflagellated, green alga *Chlamydomonas reinhardtii* (Fig. 1a, b), for which a large molecular toolkit is available (9–11), has become a model for studying the molecular interactions between microalgae and microbes (7). *C. reinhardtii* occurs mainly in wet soil ecosystems (7) and can establish mutualistic carbon-nitrogen metabolic exchange mechanisms with fungi (12) or bacteria such as *Methylobacterium* spp. (13). Moreover, algal-bacterial consortia have been used to mutualistically enhance natural hydrogen production of *C. reinhardtii* (14). However, *C. reinhardtii* can also be prone to be attacked by antagonistic bacteria. For example, the soil bacterium *Streptomyces iranensis* releases the algicide azalomycin F, which is toxic for *C. reinhardtii* unless the alga protects itself among the mycelia of the fungus *Aspergillus nidulans* (15). In a previous study, we showed that another soil bacterium, *Pseudomonas protegens* Pf-5, known to produce a wide variety of secondary metabolites (16), can inhibit the growth of *C. reinhardtii* (17). Specifically, *P. protegens* Pf-5 releases the cyclic lipopeptide orfamide A that causes a spike in cytosolic Ca^2+^ and rapid loss of flagella/cilia. However, co-cultures using an orfamide A null-mutant of *P. protegens* Pf-5 still inhibited *C. reinhardtii* growth for a minimum of eight days (17), leading us to hypothesize that at least another bacterial secondary metabolite might be involved in the antagonism of *P. protegens* Pf-5 on *C. reinhardtii*. Here, we report the discovery of a novel and unusual bacterial polyyne, protegencin, that plays a key role in the algicidal activity of *P. protegens* Pf-5: it destroys the eyespot, a primitive visual system (18, 19) (Fig. 1a, b) and causes lysis of the algal cells.

**Fig. 1:**
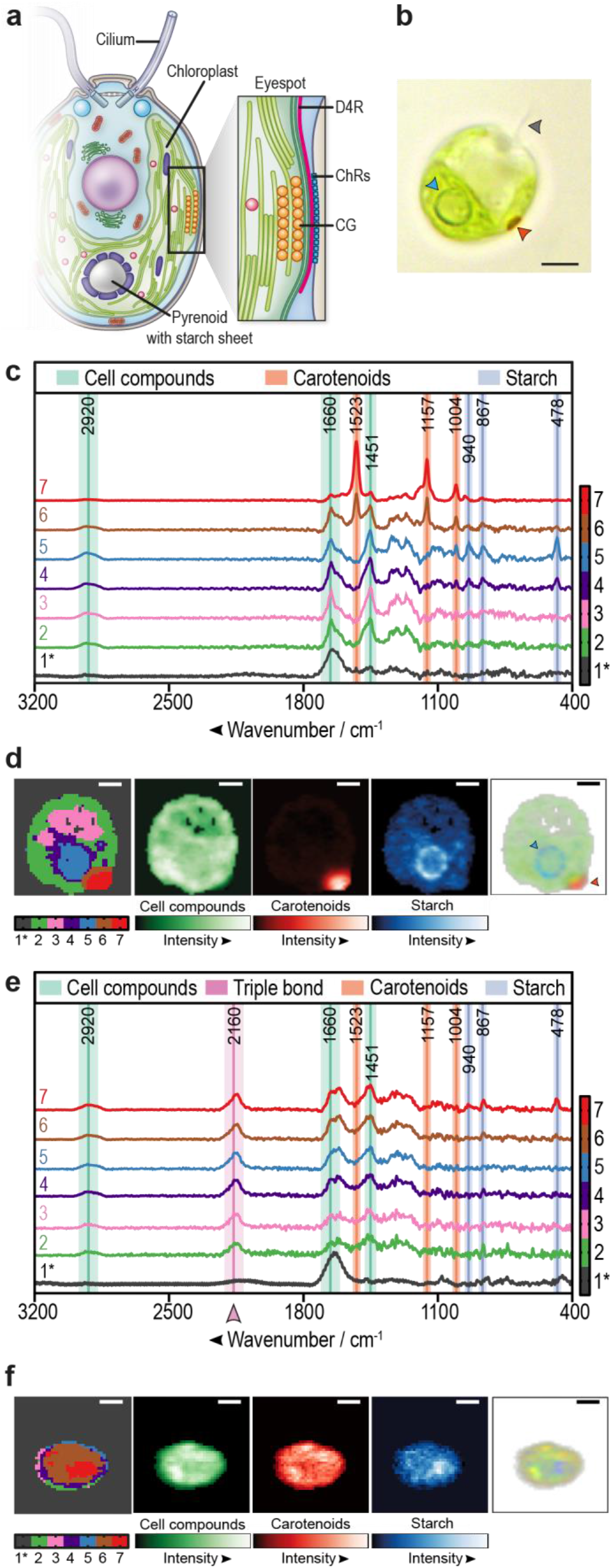
Raman microspectroscopy of *C. reinhardtii* highlights the loss of its eyespot and a novel compound in bacterial co-culture. **a**, Simplified scheme of *C. reinhardtii* (modified from (7)) with enlarged eyespot area (19). CG, carotenoid-rich lipid globules; D4R, D4 rootlet; ChRs, channelrhodopsins. **b**, Brightfield microscopy of *C. reinhardtii*. Arrowheads highlight the cilium (grey), pyrenoid (blue) and eyespot (orange). **c**, Raman spectroscopy of a representative *C. reinhardtii* cell grown in mono-culture. An algal overnight culture in TAP was fixed with 4% formaldehyde, embedded in 0.5% TAP agarose for single cell analysis. The color-coded Raman spectra display the individual cluster spectra found by k-means cluster analysis of a typical algal cell. Raman marker bands are labeled green (cell compounds), blue (starch), and orange (carotenoids). Note that phenylalanine and carotenoids share a marker band at 1,004 cm^−1^ (Supplemental Table 1). The black spectrum (1*) represents the vector-normalized background generated from areas with low content of detectable biological material. All other displayed cluster spectra are background corrected and represent clusters of components rich in cell compounds (spectra 2–7), starch (spectra 4, 5), or carotenoids (spectra 6, 7). **d**, Color-coded spatial distribution of Raman spectral components of a representative *C. reinhardtii* cell (left) as shown in **c**. Green, orange, and blue false color maps represent Raman intensities of cell compounds, carotenoids, and starch, respectively. The composite RGB image (right) is created by overlaying the normalized sums of the marker band regions associated with cell compounds (green), starch (blue), or carotenoids (red). The opacity in each pixel is proportional to the overall intensity in each pixel. Arrowheads highlight the starch sheets around the pyrenoid (blue) and the eyespot (orange). **e**, Raman spectra of a representative *C. reinhardtii* cell after overnight co-cultivation with *P. protegens* (algae:bacteria 1:1000) in TAP medium. See **c** for further details. **f**, Spatial distribution of the Raman spectroscopic clusters, integrated intensities and composite RGB image of a representative *C. reinhardtii* cell after overnight co-cultivation with *P. protegens*. See **d** for details. **b, d, f**, Scale bars: 3 μm. **c–f**, Exemplary cells were taken from the 16 h series (Fig. 2a).

## Results

### Raman microspectroscopy detects changes in algal eyespot carotenoids and a new compound in algal-bacterial co-cultures

To understand the antagonistic interplay between *C. reinhardtii* and *P. protegens* Pf-5 (thereafter abbreviated as *P. protegens*), we aimed to spatially resolve the molecular composition of microalgal samples in absence and presence of the bacteria with subcellular resolution using Raman microspectroscopy (20). Due to the non-invasive nature of Raman microspectroscopy and the fact that water only mildly distorts Raman spectra, this technique has been well suited to study biological samples (21) such as bacteria and microalgae (22–28), including *C. reinhardtii* (26–28).

Axenic algal cultures were first analyzed to establish a reference baseline for comparisons. To obtain efficient hyperspectral Raman images, *C. reinhardtii* cells were fixed in 4% formalin and embedded in 0.5% agarose to immobilize the cells (Supplementary Fig. 1); cells embedded solely in agarose were not sufficient to immobilize the algae and were therefore not suitable for spatially-resolved subcellular imaging. Raman images were recorded in the spectral range from 104 cm^−1^ to 3,765 cm^−1^ using an excitation wavelength of 785 nm. Raman bands were assigned to relevant compounds according to characteristic wavenumbers listed in Supplementary Table 1. The spectral regions from 400 to 1,750 cm^−1^ and 2,800 to 3,200 cm^−1^ include typical marker bands for cell compounds (lipids and proteins), carotenoids, and starch (Fig. 1c). Raman intensity maps were calculated by the sum of intensities over specific compound marker bands in a spatially-resolved manner (Fig. 1d and Supplementary Fig. 2). Two sub-organelles were especially well visualized by Raman microspectroscopy: (i) the eyespot, based on its enriched carotenoids, and (ii) the pyrenoid, situated in the U-shaped chloroplast and detectable by its surrounding starch sheet layer (Fig. 1a, d). It should be noted that the three-dimensional image of the cell within the agarose bed (Fig. 1) is projected on to two-dimensions and thus the eyespot often appears to be localized non-horizontally.

We then analyzed *C. reinhardtii* cells that were grown in co-culture overnight with *P. protegens* Pf-5. Here, two major Raman spectroscopic changes were observed: (i) algal cells showed strongly reduced peaks at wavenumbers 1,523 cm^−1^, 1,157 cm^−1^, and 1,004 cm^−1^, which are all assigned to carotenoids (Fig. 1e, Supplementary Table 1), and (ii) a new peak appeared at ~2,160 cm^−1^, which lies in the so-called “silent wavenumber region” for which there is no known overlap with Raman signatures of biological origin. This 2,160 cm^−1^ peak indicates the presence of a triple-bond containing compound (29). Interestingly, many *C. reinhardtii* cells examined in co-culture with *P. protegens* had lost their typical eyespot (Fig. 1f). To determine whether *P. protegens* produces the compound responsible for the peak in the silent wavenumber region, we performed Raman microspectroscopy on axenic bacterial cultures, which also showed a peak at ~2,150 cm^−1^ (Supplementary Fig. 3).

To study the appearance of the novel triple-bond-bearing compound and the loss of the eyespot in bacterial-algal co-cultures in greater detail, we incubated the *C. reinhardtii* cells with *P. protegens* for 16 h and 24 h, using axenic algal cultures cultivated for the same periods of time as reference controls. In the Fig. 2a, the carotenoid clusters are highlighted at 1,523 cm^−1^, 1,157 cm^−1^, and 1,004 cm^−1^, although a typical protein marker band for phenylalanine is also detected at 1,004 cm^−1^ (30). The intensity ratio between 2,160 cm^−1^ and 1,451 cm^−1^ (C-H-deformation mode, the most prominent cell compound feature) was calculated as a measure for the presence of the triple-bond compound for each algal cell (Fig. 2b). These experiments confirmed that axenic algal cultures did not contain the novel substance, and that it was only observed in co-culture of *C. reinhardtii* with *P. protegens* (Fig. 2a, b). Moreover, we observed a clear reduction in the number of carotenoid clusters in the presence of *P. protegens* (Fig. 2b). After 16 h in co-culture, some algal cells appeared enlarged and eyespots often disintegrated into smaller size particles or were found to be absent as observed by brightfield microscopy (Fig. 2c, g). After 24 h in co-culture, algal cells appeared to lyse and in most cells eyespots were completely undetectable (Fig. 2c, g). Visualizing the spatial distribution of the triple-bond compound in algal cells co-cultured with *P. protegens* for at least 16 h, we found that the substance sometimes accumulates in local spots but is also distributed throughout *C. reinhardtii* (Supplementary Fig. 4).

**Fig. 2:**
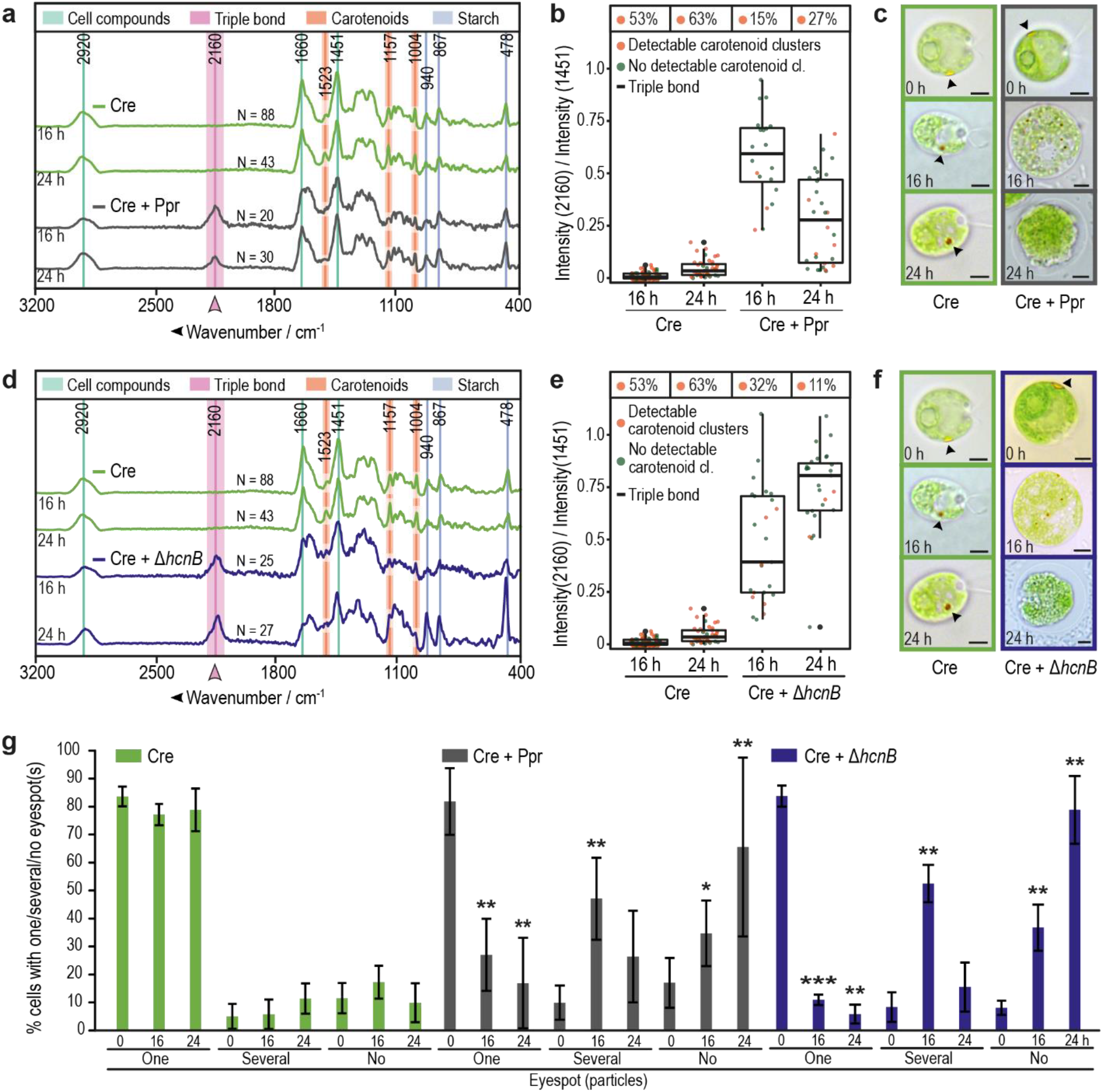
Disappearance of eyespot carotenoids coincides with the appearance of a triple bond-containing compound (TC). **a, d**, Average Raman spectra of *C. reinhardtii* show a peak at 2160 cm^−1^ and reduction of detectable carotenoid peaks over time when co-cultured with *P. protegens* (**a**, Cre + Ppr) or its Δ*hcnB* mutant (**d**, Cre + Δ*hcnB*) compared to axenic *C. reinhardtii* (**a, d**, Cre). Cultures were grown for 16 or 24 h in TAP. Spectra of algal cells were generated by averaging over all pixels for the relevant Raman spectral class within the cell body. N = total number of analyzed cells. All data were obtained from at least three independent experiments. **b, e**, Summary of Raman spectroscopic measurements from **a, d** show the presence of the TC (box plot) and a reduction of carotenoid clusters in co-cultures (Cre + Ppr) and (Cre + *ΔhcnB*). The data points plotted are for the ratio of average Raman intensities associated with the TC (2,160 cm^−1^) vs. a dominant band representing *C. reinhardtii* cellular material (C-H deformation mode at 1,451 cm^−1^). Each point represents a measurement of a single alga; box plots indicate quartiles; % values above each box summarize the fraction of cells with detected carotenoid cluster(s). **c, f**, Representative brightfield microscopic images of axenic *C. reinhardtii* (green) and in co-culture (**c**, Cre + Ppr, grey, **f**, Cre + *ΔhcnB,* blue) after 0, 16, or 24 h show the disappearance of eyespots (see arrowhead) over time in the presence of bacteria. Scale bars: 3 μm. **g**, Fraction of algal cells (from **a–f)** with eyespots as determined by brightfield microscopy. After 16 h in co-culture, the eyespot is mostly disintegrated, and undetectable after 24 h, whereas most cells in the axenic culture maintain one eyespot. Asterisks indicate significant differences as calculated by the Kruskal-Wallis test with Dunn’s post-hoc test (*, P < 0.05; **, P < 0.01; and ***, P < 0.001) in co-culture compared to axenic *C. reinhardtii* at the same time point. Error bars indicate standard deviations with N ≥ 300 cells per time point and culture.

We surmised that one possible triple-bond containing compound responsible for the peak in co-culture could be hydrogen cyanide, which is known to be produced by *P. protegens* (16). Using the Δ*hcnB* cyanide mutant JL4809 (16), we determined whether the carbon-nitrogen triple bond of cyanide is responsible for the 2,160 cm^−1^ peak. This peak was clearly detectable after 16 h and 24 h in Δ*hcnB* mutant-*C. reinhardtii* co-cultures (Fig. 2d, e). The disappearance of the eyespot carotenoid clusters was still observed in many cells (Fig. 2e), and brightfield micrographs of co-cultured algal cells looked identical to those co-cultured with wild-type *P. protegens* (Fig. 2c, f and g). These data suggest that hydrogen cyanide is not the relevant triple-bond compound.

### Identification of the polyyne protegencin by analytical chemistry and generation of a protegencin null mutant

Next, we considered polyynes, fatty acid derivatives with multiple C-C triple bonds, as alternatives. In bacteria, polyyne biosynthesis is encoded in gene clusters bearing genes for desaturases, a fatty acyl-AMP ligase, an acyl carrier protein, and a thioesterase (31). Mining the genome (32) of *P. protegens* revealed a tentative polyyne biosynthesis gene cluster *(pgn*) for a compound we name “protegencin”. According to homology searches, *pgnE*, *pgnF,* and *pgnH* encode desaturases and a thioesterase, *pgnD* encodes a fatty acyl-AMP ligase, and *pgnG* encodes an acyl carrier protein (Fig. 3a). The predicted gene cluster includes genes PFL_0258 to PFL_0268, a previously defined orphan gene cluster with an unknown metabolic product (33). It is an orthologous biosynthetic gene cluster to the polyyne caryoynencin (YP_257407 – 257414 (31)). Parts of this gene cluster (PFL_0261–0267) are also homologous to a gene cluster for the biosynthesis of the collimonins, polyynes produced by *Collimonas fungivorans* (34, 35). We predicted that *P. protegens* would produce at least one polyyne compound from the *pgn* gene cluster that could account for the peak at 2,160 cm^−1^.

**Fig. 3:**
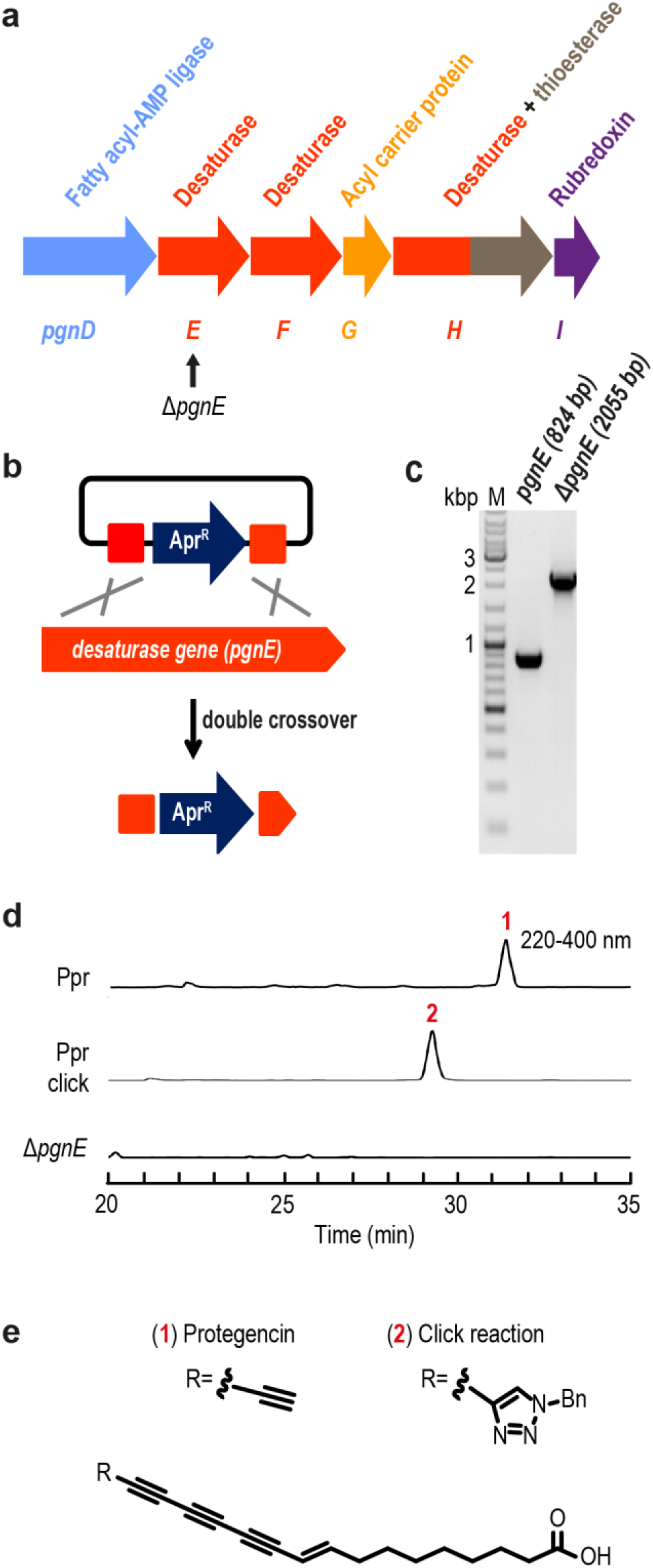
Creation of a *P. protegens* mutant deficient in protegencin production and chemical characterization of protegencin. **a**, *Protegencin* (*pgn*) biosynthesis gene cluster (PFL_0262–0267) with knock-out target gene, *pgnE,* indicated (PFL_0263, black arrow). **b**, Double crossover strategy used to obtain a *P. protegens* mutant deficient in protegencin production. **c**, Agarose gel shows successful insertion of the *Apr* gene in *pgnE*. **d**, HPLC analysis (PDA 220–400 nm) of *P. protegens* culture extracts, Click reaction product, and of Δ*pgnE* mutant extracts. **e**, Structure of protegencin as determined from NMR and high resolution MS analysis (Supplementary Fig. 5–10); Bn, Benzyl. Experiments were repeated at least three times.

**Fig. 4:**
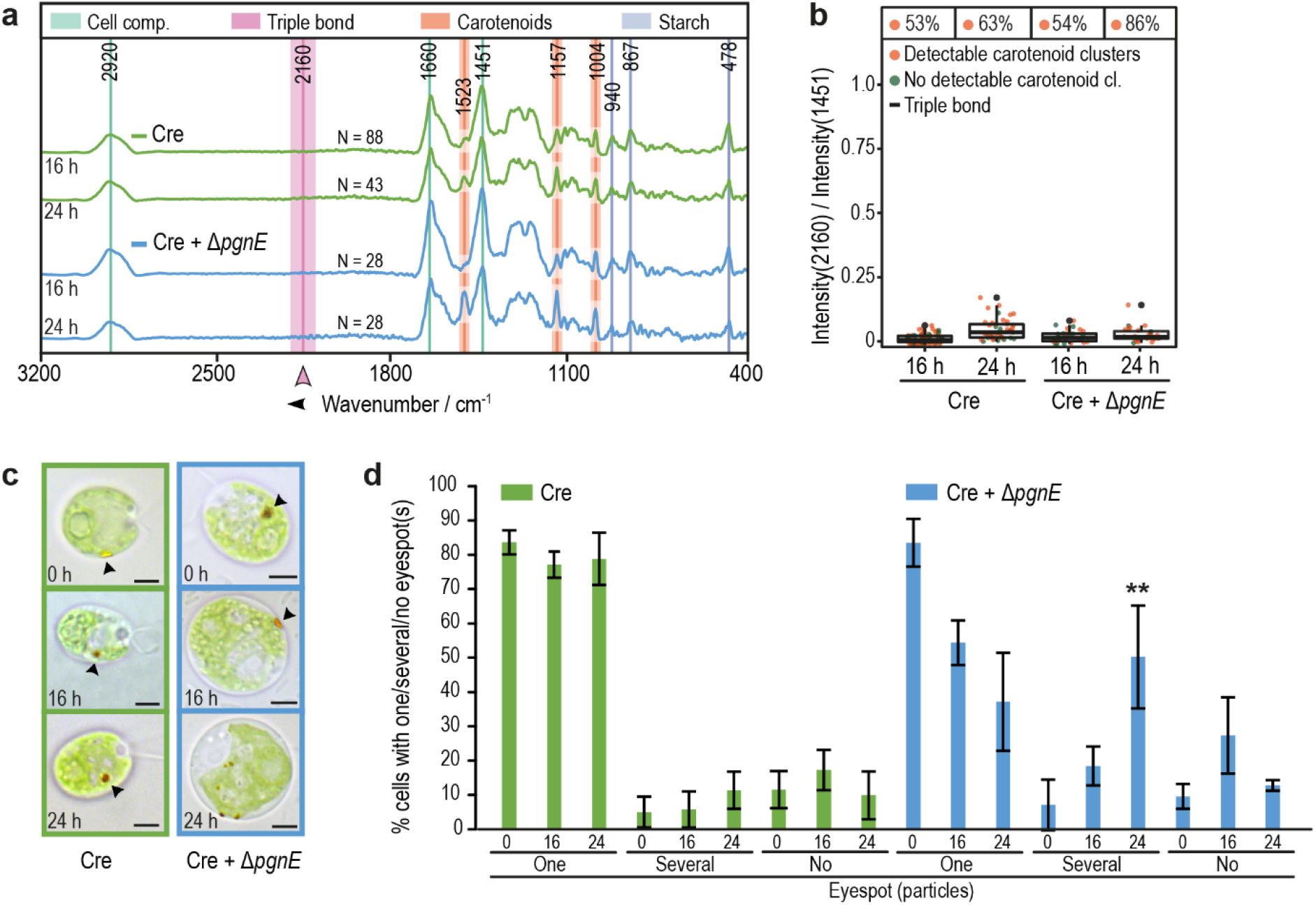
Identification of protegencin in co-cultures and its effects on carotenoid clusters. **a**, Raman spectroscopy of axenic *C. reinhardtii* (green, Cre) and in co-culture with the *P. protegens pgnE* null mutant (Cre + Δ*pgnE*, light blue) suggests that protegencin is the compound responsible for the peak at 2,160 cm^−1^, which remains absent even after 24 h of co-culturing (N = total number of analyzed cells). All data were obtained from at least three independent experiments. See legend of Fig. 2a for further details. **b**, Summary of Raman spectroscopic measurements presented in **a**, with % above each box representing the total fractions of algal cells with carotenoid cluster(s). See legend of Fig. 2b for further details. **c**, Representative brightfield micrographs of axenic *C. reinhardtii* (green) and in co-culture with the *P. protegens* Δ*pgnE* null mutant (blue) after 0, 16, or 24 h show that the eyespot disintegrates slower than in co-cultures with *P. protegens* wild type or *ΔhcnB* mutant strains (see Fig. 2c, f; arrowheads highlight the eyespot). Scale bars: 3 μm. **d**, Fraction of algal cells with eyespots as determined by brightfield microscopy of cultures analyzed in **a-c** after 0, 16 and 24 h. After 16 h in co-culture, the eyespot is mostly still present; after 24 h, it disintegrates in more than one eyespot in about half of the cells. Asterisks indicate significant differences as calculated by the Kruskal-Wallis test with Dunn’s post-hoc test (*, P < 0.05; **, P < 0.01; and ***, P < 0.001) in co-culture compared to axenic *C. reinhardtii* at the same time point. Error bars indicate standard deviations with N ≥ 300 cells per time point and culture.

To test if protegencin accounts for the 2,160 cm^−1^ Raman peak, we created a targeted *pgn* null mutant (Δ*pgnE*). Specifically, we inactivated the desaturase gene, *pgnE*, by means of a double-crossover knock-in strategy involving the introduction of an apramycin resistance gene (Apr^R^) at the *pgnE* locus (see Methods, Fig. 3a–c).

By comparative HPLC analyses (PDA 220–400 nm) of ethyl acetate extracts of wild-type and Δ*pgnE P. protegens* cultures (Fig. 3d, Supplementary Fig. 5), we detected a candidate polyyne compound with a characteristic UV signature. We succeeded in the isolation of this compound, protegencin, and elucidated its structure using high resolution MS and NMR spectroscopic analysis (Fig. 3e, Supplementary Fig. 5–10, Supplementary Table S2). Protegencin is a highly unsaturated octadecanoid fatty acid with four conjugated triple bonds and one double bond. A copper-catalyzed alkyne-azide cycloaddition (CuAAC) reaction with benzyl azide confirmed that this compound has a reactive terminal triple bond (Fig. 3d, e and Supplementary Fig. 5).

Metabolic profiling of the verified Δ*pgnE* mutant (Fig. 3c) showed that protegencin production was completely abolished (Fig. 3d).

### Protegencin causes the disappearance of eyespot carotenoids in co-culture

In *C. reinhardtii* co-cultures with the Δ*pgnE* mutant, the 2,160 cm^−1^ Raman peak was absent (Fig. 3a, b), consistent with the lack of expressed protegencin. We confirmed that protegencin is responsible for the 2,160 cm^−1^ Raman peak by predicting its Raman spectrum by means of density functional theory (DFT). DFT calculations predicted characteristic features between 2,100 cm^−1^ and 2,200 cm^−1^, in good agreement with empirically observed peak features (Supplementary Fig. 3).

We observed that eyespot carotenoid clusters in *C. reinhardtii* were not affected by co-culture with the Δ*pgnE* mutant strain (Fig. 3b). These findings suggest that protegencin is the cause for the disappearance of the algal eyespot carotenoid clusters. Consistent with this, after 16 h in co-culture with the Δ*pgnE* mutant, a single eyespot was visible in most *C. reinhardtii* cells by brightfield microscopy; after 24 h in co-culture some algal cells had a single eyespot while others had more than one eyespot (Fig. 3c, d). In the absence of protegencin expression, *C. reinhardtii* eyespots are not degraded in co-culture with *P. protegens*.

### Protegencin lyses and kills algal cells

To corroborate the role of protegencin, we evaluated the direct effect of purified compound on *C. reinhardtii* cells, focusing on the integrity of the algal cell membrane as assayed by Evans blue exclusion. Mastoporan, a toxin from wasp venom, known to cause lysis and death of *C. reinhardtii*, was used as a positive control at a concentration of 10 μM (36). In the presence of mastoporan, about 90% of the cells of *C. reinhardtii* were stained with Evans blue after 30 sec (Fig. 5a). Protegencin, which was dissolved in DMSO after its purification (see Supplementary Methods), was used at a similar concentration (2% v/v equivalent to 10.4 μM), as determined by the click reaction and HPLC analysis (see Methods). Protegencin treatment resulted in Evans blue staining of >95% of the cells after 24 h of incubation. This experiment was repeated with a lower concentration of protegencin (0.5% v/v), which had the same effect (Fig. 5a). These data show that protegencin can efficiently perforate *C. reinhardtii* cell membranes even at low micromolar concentrations. We also evaluated the toxicity of protegencin on *C. reinhardtii* after short incubation with 0.5 % v/v protegencin by survival assays (modified from (37)). Incubation with protegencin for only 1 h reduced colony forming units (CFUs; visible after 7 days) down to about 10% (Fig. 5b); an incubation time of 4 h resulted in no CFUs (Fig. 5b, right panel), supporting the highly toxic effect of protegencin on *C. reinhardtii*.

**Fig. 5:**
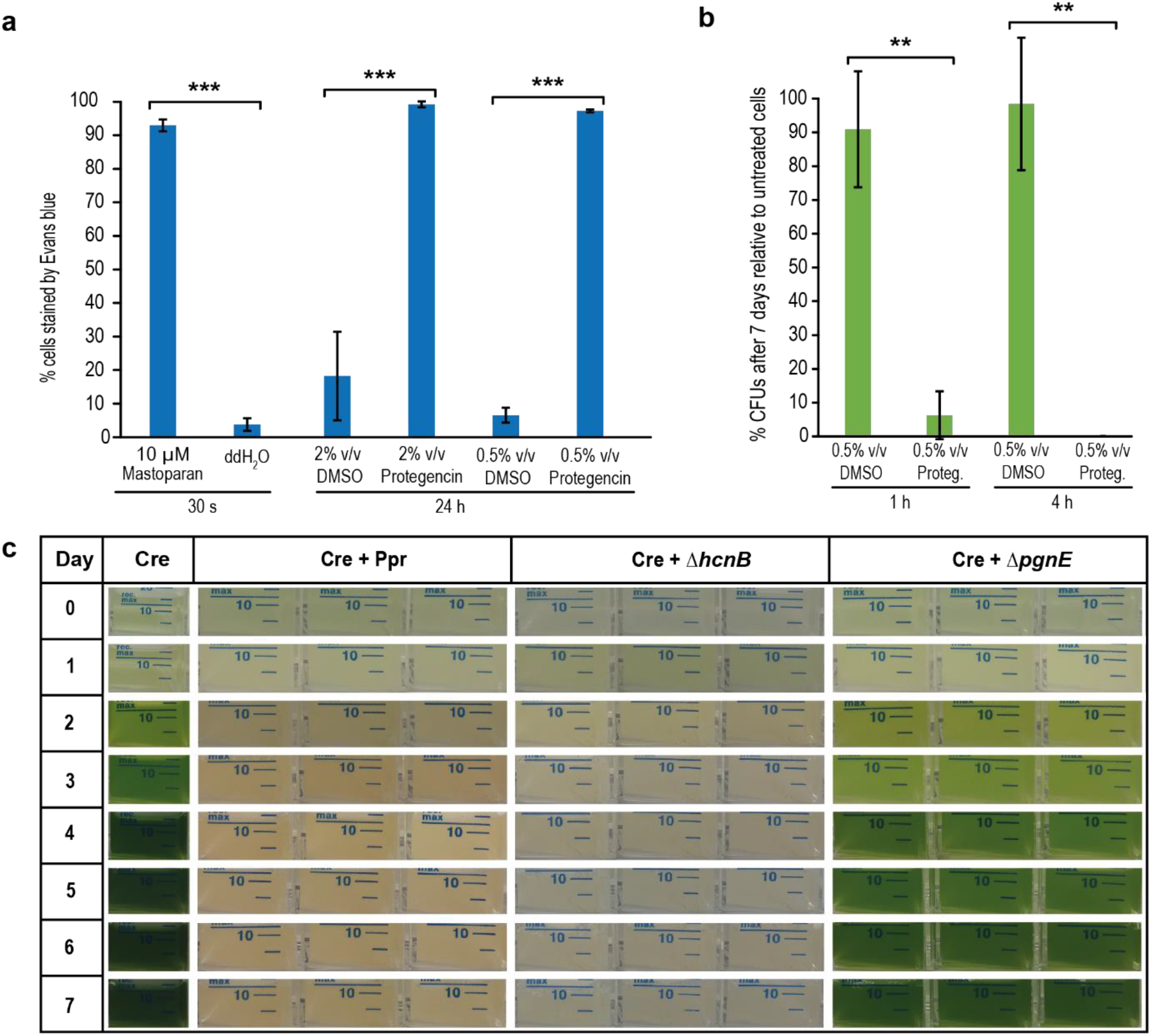
Protegencin, a potent toxin, lyses and bleaches algal cells. **a**, Protegencin effectively damages the cell membrane of *C. reinhardtii* after 24 h of incubation as evaluated by Evans blue staining. Mastoparan was used as positive control. As negative controls, DMSO (solvent for protegencin) and water (solvent for mastoparan) were used. For each treatment, three independent replicates with three technical replicates each were examined; for each technical replicate per time point and treatment, N ≥ 500 cells were analyzed. **b**, Within 4 h of incubation with 0.5 % (v/v) purified protegencin, *C. reinhardtii* CFU growth was prevented (over 7 days). Control, DMSO. For each treatment, three independent replicates with three technical replicates each were examined. Error bars indicate standard deviations and asterisks indicate significant differences compared to Cre, calculated by Student’s t-test (**, P < 0.01; and ***, P < 0.001) **c**, In liquid co-cultures of *C. reinhardtii* with *P. protegens*, Δ*hcnB*, or Δ*pgnE* mutants, algal growth is only possible when protegencin production is prevented. Experiments were conducted three times independently, each with three technical replicates.

To test whether protegencin is the primary metabolite causing growth arrest and bleaching of algal cells, we compared co-culture growth of *C. reinhardtii* with *P. protegens* wild type, Δ*hcnB* cyanide mutant, or Δ*pgnE* mutant strains over a period of seven days. Axenic *C. reinhardtii* cultures were used as controls (Fig. 5c). As observed previously (17), *C. reinhardtii* cells do not grow in co-culture with *P. protegens*. The characteristic chlorophyll-based green color of algal mono-cultures is absent in co-cultures due to growth arrest (Fig. 5c, (17)). In co-culture with the Δ*hcnB* cyanide mutant, no algal growth is visible either. However, the algal cells grow well in co-culture with the Δ*pgnE* protegencin mutant (Fig. 5c, right panel). These findings corroborate that protegencin is the major toxin secreted by *P. protegens* against *C. reinhardtii*.

## Discussion

Although microalgae are key contributors to global carbon fixation and fundamental to food webs, their interaction with other microbes is still not well understood at a molecular level. Recently, several studies on algicidal bacteria and their natural products that affect the performance of microalgae have been reported (reviewed in (3)), and have provided the first insights into the mechanisms of algicidal bacteria. However, studies involving *in situ* chemical analysis of microalgal cells or genetic manipulations to alter the production of the bacterial secondary metabolites and nature of the microbial interaction are still rare.

Here, we identified and characterized a natural product underlying an antagonistic interaction between the green microalga *C. reinhardtii* and the bacterium *P. protegens,* both model soil organisms, under laboratory conditions. Using genetically constructed mutants, brightfield microscopy, label-free Raman microspectroscopy, high resolution MS, and NMR analyses, we discovered a novel polyyne bacterial toxin, protegencin, that targets algal eyespots and causes algal cell lysis.

*P. protegens* produces a variety of secondary metabolites controlled by the GacS-GacA regulatory system (16, 33, 38), including the *pgn* cluster. Nevertheless, the production of a polyyne was thus far only predicted *in silico* (31). Polyynes like protegencin are notoriously unstable compounds. They have been isolated from different organisms, including a variety of land plants, fungi, and insects. However, only few bacterial polyynes are known of ecological relevance (31, 39–41). The first identified bacterial gene cluster coding for the biosynthesis of a polyyne was caryoynencin from *Burkholderia* spp. (31). In a former genome mining analysis, orthologous *pgn* gene clusters were also found in *Collimonas* and *Mycobacterium* as well as in different *Burkholderia* species (31). Currently, it is not known whether these bacteria can also exert algicidal activities using polyynes and interactions of these bacteria have only been studied with organisms other than photosynthetic protists. Some bacterial polyynes have been shown to protect other organisms; for instance, the polyyne caryoynencin is part of a blend of antibiotics produced by symbiotic *Burkholderia gladioli* bacteria to protect the egg stage of a group of herbivorous beetles against detrimental bacteria (42). Cepacin from *Burkholderia ambifaria* exerts biocontrol in *Pisum sativum* by protecting the crop from an attack of oomycetes (43). Cepacin A and B, isolated from *Pseudomonas cepacia*, exert activities against other microbes, e.g. staphylococci (44). The polyyne collimomycin, later renamed to collimonin, from the bacterium *Collimonas fungivorans* inhibits the growth of *Aspergillus niger* hyphae and controls hyphal branching and pigmentation (34, 35). In contrast, protegencin characterized in this study plays an unprecedented role as a potent bacterial toxin against the green microalga *C. reinhardtii*.

It will be intriguing to study in the future if other microalgae than *C. reinhardtii*, living in freshwater and soil, such as *Gonium pectorale*, *Haematococcus pluvialis* (both Chlorophyta) or *Euglena gracilis* (Euglenophyta), are also affected by protegencin. Previously, we have shown that another *P. protegens* natural product, orfamide A, immobilizes select Chlorophyte algae (17). Orfamide A triggers an increase in cytosolic Ca^2+^ but is unable to penetrate the algal cells even at a concentration of 35 μM within 30 sec (17). In contrast, protegencin permeates *C. reinhardtii* cells. These toxins from *P. protegens* are not only chemically different but also functionally distinct. Considering the timescale for the action of the two compounds, these bacteria may use a two-step strategy to antagonize algae: (i) quick immobilization (deflagellation) of algal cells by orfamide A and thereafter (ii) an artillery-like attack using polyynes like protegencin whereby they “blind” them by destroying the eyespot and lyse them (Fig. 6).

**Fig. 6:**
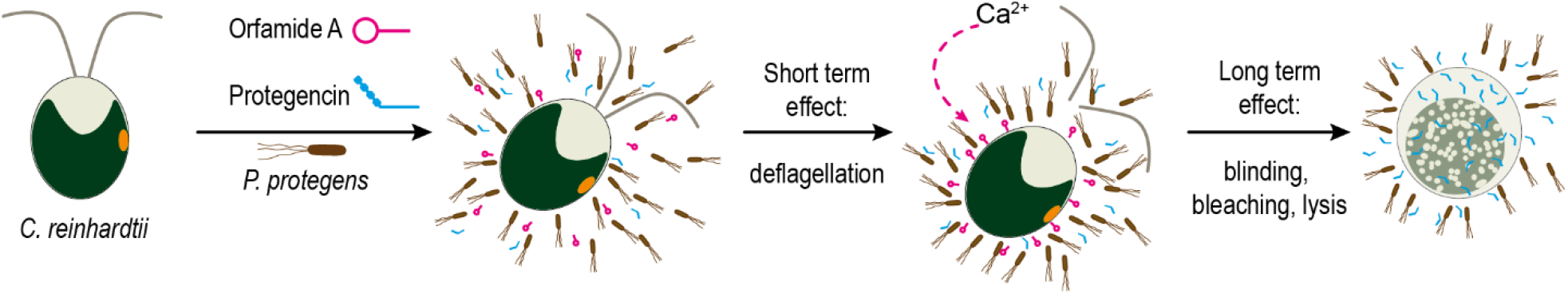
Proposed scheme of *P. protegens’* algicidal activity towards *C. reinhardtii*. Harmful secondary metabolites orfamide A (magenta) and protegencin (cyan) are produced by *P. protegens*. Orfamide A triggers an increase in cytosolic Ca^2+^ that results in deflagellation of the algae within minutes (17). Slower (on the order of hours), more potent damage by protegencin resulting in eyespot decay (“blinding”), bleaching, and algal cell lysis occurs subsequently.

In their natural habitats, microalgae interact with other microbes via a variety of chemical mediators, resulting in mutualistic or antagonistic relationships. This study sheds light on the chemical weapons used by antagonistic bacteria and their mode of action in microalgal communities. It reveals fundamental mechanisms for how fitness of photosynthetic microbes can be influenced, and specifically, the role and mode of action of polyynes in the arsenal of algicidal bacteria.

## METHODS

### Strains and culture conditions

*C. reinhardtii* strain SAG 73.72 (mt^+^) that was obtained from the algal culture collection in Göttingen, was used as wild type. *C. reinhardtii* was grown in liquid TAP (Tris-Acetate-Phosphate) medium (45) at 23 °C under a 12:12 light-dark (LD) cycle with a light intensity of 55 μE and constant orbital shaking (100 rpm). *P. protegens* strain Pf-5 was used as bacterial wild type (46). A cyanide mutant, Δ*hcnB* mutant, JL4809 (16), was used as well as a Pf-5 strain deficient in protegencin production (see below), Δ*pgnE*. Unless stated otherwise, axenic bacteria were grown in LB medium at 28 °C with constant orbital shaking (200 rpm).

### Algal bacterial co-cultures

For the co-cultures, *C. reinhardtii* cells were precultured to a cell density of 3–6 × 10^6^ cells mL^−1^. Bacterial pre-cultures were grown to an optical density OD_600_ > 4 (as determined by serial dilutions of the bacterial pre-cultures) overnight. Prior to co-cultivation, bacterial cells were washed twice with TAP medium. All cells were then reinoculated in TAP medium with a starting algal cell density of 2 × 10^5^ cells mL^−1^ and a bacterial cell density of 2 × 10^8^ cells mL^−1^. The co-cultures were grown under the conditions mentioned above for the growth of *C. reinhardtii*. As control, axenic *C. reinhardtii* cultures were inoculated at a starting cell density of 2 × 10^5^ cells mL^−1^ and grown and analyzed accordingly.

### Creation of a *P. protegens* protegencin mutant

To inactivate the gene *pgnE* of the protegencin gene cluster *P. protegens*, a target double crossover strategy was chosen to insert the apramycin resistance gene from PIJ773 into each cay gene. Details are presented in Supplemental Methods.

### Sample preparation for Raman imaging

Axenic *C. reinhardtii* and co-cultures of *C. reinhardtii* with *P. protegens* wild type and mutants were grown as mentioned before. 1.5 mL cell suspension were removed under sterile conditions, fixed in 4% (v/v) formalin for 10 min and centrifuged for 5 min at 4,500 × *g*. Afterwards, the supernatant was removed, and the cell pellet was re-suspended in 375 μL TAP. The obtained suspension was mixed with 375 μL hand warm 1% (w/v) agarose in TAP. The agarose-cell mixture was quickly transferred onto CaF_2_ substrate and cooled down at 4 °C for at least 15 min. The sample substrates were covered with distilled water before Raman measurements.

### Hyperspectral Raman imaging

Hyperspectral Raman images were acquired using a confocal Raman imaging microscope (alpha 300 R, WITec, Ulm, Germany) with a water immersion objective (Nikon Corporation, Tokyo, Japan; magnification: 60×; numerical aperture: 1.0). The excitation light with a wavelength of 785 nm was provided by a cw diode laser (Toptica Photonics, Gräfelingen, Germany). The waist diameter of the confocal volume was estimated by calculating the diameter of the central Airy disc and is ~0.96 μm (d_Airy_ = 1.22 λ_Laser_ / NA_Objective_; λ_Laser_ = 785 nm, NA_Objective_ = 1.0). The excitation power measured at the front lens of the objective was ~70 mW. Raman scattered light was dispersed by a diffraction grating (300 grooves mm^−1^) and detected by a cooled EM-CCD-detector in a range from 104 cm^−1^ to 3,765 cm^−1^. Algal samples were selected under the microscope and single algae cells were centered in a 15 μm × 15 μm large x,y-scanning grid. The hyperspectral Raman data cube was generated by point-wise integration of the scanning grid in x- and y-direction with a step-size of 0.34 μm equal in both spatial directions. Two independent scans of the area incorporating a single algal cell were performed. In an initial rapid scan (integration time: 100 ms/pixel) the autofluorescence of the algae was bleached to decrease the background fluorescence level. The subsequent acquisition scan was performed with a dwell time of 500 ms/pixel and a hyperspectral data cube I (x, y, ν) with the dimension of 45 pixel × 45 pixel × 1024 wavenumbers was obtained.

### Pre-processing of the Raman spectra

All raw data was processed and analyzed by an in-house developed script in the programming language R (version 4.0.2) (47). The pre-processing is described in detail in references (48) and (49). In short, the Raman spectra were cleared from cosmic spikes and wavenumber calibration was performed with 4-acetaminophenol.

Subsequently the wavenumber region was restricted to 350 cm^−1^ – 3,300 cm^−1^. The wavenumber axis of all spectra were interpolated using a cubic fmm-spline (new wavenumber pitch: 3 cm^−1^) and background corrected by employing a statistic-sensitive-non-linear iterative peak-clipping algorithm (smoothing = true, iteration = 100, window = “3”) (50). To minimize the influence of the agarose embedding on Raman algal cell spectra, an additional correction was performed accounting for the spectral influence of the agarose (see supplementary information for computational details).

## Supporting information

Supplemental Material to Hotter et al.

## Acknowledgements

We thank Erik Hom for proof-reading the manuscript, Georg Kreimer for helpful comments on Fig. 1a and Debbie Maizels for its design. Our work was supported by fellowships of the International Leibniz Research School ILRS (under the head of the Jena School for Microbial Communication) awarded to V.H. and P.A. D.Z., A.S., S.S., C.H., J.P. and M.M. were funded by the Deutsche Forschungsgemeinschaft (DFG; German Research Foundation) by SFB 1127/2 ChemBioSys - 239748522.

## Author Contributions

J.P., C.H., and M.M. designed research; V.H., D.Z., H.J.K., A.S., J. F., J. H., C.M. and P.A. performed research. V.H., D.Z., J.H., H.J.K., A.S., C.M., P.A, M.S., S.S., J.P., C.H., and M.M. analyzed data. S.S. contributed to Fig. 1A. J.L. and Q.Y. provided the *hcnB* mutant. V.H., D.Z., H.J.K., C.H. and M.M. wrote the paper, with input from all co-authors.

## Correspondence

Correspondence should be addressed to MM.

