## Supplemental Material to Hotter et al. for "A polyyne toxin produced by an antagonistic bacterium blinds and lyses a green microalga"

### Supplementary information

**This PDF file includes:**

Supplementary Materials and Methods

Supplementary Figures 1 to 10

Supplementary Tables 1 to 3

Supplementary References

#### Creation of a desaturase mutant (Δ*pgnE*)

*P. protegens* Pf-5 (thereafter abbreviated as *P. protegens*) was cultivated on LB agar at 30 °C for 2 days and inoculated into Luria-Bertani (LB) medium for further incubation for 1 day. The bacterial cells were collected by centrifugation (5,000 rpm, 5 min). Genomic DNA of *P. protegens* was acquired by using a MasterPure$^{TM}$ complete DNA purification kit (Lucigen, Madison). The *pgnE* gene in the putative protegencin biosynthetic gene cluster was PCR amplified from the genomic DNA of *P. protegens* with specific primers (Supplementary Table 3) and ligated with the apramycin resistance gene. The combined gene fragment was inserted into pJET1.2/Blunt vector. *Escherichia coli* TOP10 competent cells were transformed with the corresponding vector using electroporation (2,500 V). The transformed cells were incubated on LB agar and medium with apramycin and ampicillin at 37 °C for 1 day. The vector containing the combined gene fragment was purified by using a Monarch® Plasmid Miniprep Kit (New England BioLabs, Frankfurt am Main). The transformation of *P. protegens* with the corresponding vector was conducted by electroporation at 2,500 V, and the transformed cells were incubated on LB agar containing apramycin and ampicillin at 30 °C. The colony containing the corresponding mutation was inoculated into LB medium with apramycin and ampicillin and further incubated at 30 °C until $OD_{600}$ value became 4–5. The bacterial cells were collected by centrifugation (5,000 rpm, 5 min) and washed with Tris-Acetate-Phosphate (TAP) medium twice. The collected cells were incubated in TAP medium with ampicillin and apramycin at 30 °C for 1 day for further analytic procedures.

**General analytic procedures**

Analytic high-performance liquid chromatography (HPLC) measurements were performed on a Shimadzu Prominence HPLC system consisting of an autosampler, high-pressure pumps, column oven and PDA using a Macherey-Nagel C18 reverse phase column (Nucleosil 100, 5 µm, 125 × 4.6 mm, flow rate 1 mL $min^{-1}$). HPLC grade acetonitrile and deionized water with 0.1% trifluoroacetic acid (TFA) were used as mobile phase for HPLC. The gradient elution for HPLC operation was from 0.5/99.5 of $CH_3CN/H_2O$ with 0.1% (v/v) TFA to 100/0 for 5–35 min. Preparative HPLC measurement were performed on a Gilson Abimed equipped with Binary Pump 321 and 156 UV/Vis detector (eluents: water with 0.1% (v/v) TFA and acetonitrile) using a Macherey-Nagel C18 reverse phase column (Nucleosil 100, 5 µm, 250 × 10 mm, flow rate 5 mL $min^{-1}$). Liquid chromatography-mass spectrometry (LC-MS) measurements were performed using an QExactive Orbitrap High Performance Benchtop LC-MS with an electrospray ion source and an Accela HPLC system (Thermo Fisher Scientific, Bremen). For MS/MS measurements, an Exactive Orbitrap mass spectrometer with an electrospray ion source (Thermo Fisher Scientific, Bremen) was used. NMR spectra were recorded on Bruker AVANCE III 500 and 600 MHz (equipped with a Bruker Cryo platform) instruments.

**Protegencin isolation and CuAAC reaction**

*P. protegens* was cultivated on LB agar at 30 °C for 2 days and inoculated into TAP medium. The bacterial culture was incubated at 30 °C with orbital shaking (150 rpm) for 1 day. The bacterial culture was extracted with ethyl acetate three times. The combined organic layers were reduced under rotary evaporation. The crude extract was purified by putative HPLC and silica gel column (Hex/EtOAc = 1:1) to remove extra fatty acids. The CuAAc reaction procedure was followed as previously reported (1). NMR and MS data of the triazol compound from click reaction with protegencin and benzyl azide are same as reported one (1).

**Bright field microscopy of co-cultures**

To determine the percentage of algal cells with one and no eyespots or multiple eyespot particles in the different co-cultures, 1 ml cell suspension was taken under sterile conditions, concentrated at 4,500×*g* for 5 min at room temperature, and resuspended in 500 µl TAP. Cells were not fixed. Prior to examination, cells were allowed to settle on the microscope slide for a few minutes at room temperature in the dark. Pictures of at least 100 cells per co-culture were taken with an Axiophot (Carl Zeiss, Germany). Each time, an axenic *C. reinhardtii* culture was examined the same way as a control. The percentage of cells with one and no eyespots or multiple eyespot particles was analyzed visually. The experiments were conducted three times independently.

**Evans blue assay**

For the Evans blue assay, the algal cell density was adjusted to $2 \times 10^6$ cells $mL^{-1}$ in TAP. Subsequently, cells were treated with 0.5% (v/v) DMSO and 0.5 % (v/v) protegencin, respectively, for 24 h. 10 µM mastoparan was used as a positive control (2). For the duration of the treatment, the cells were kept shaking in the dark at room temperature. Subsequently, the cells were split in three technical replicates per treatment, washed three times in TAP and finally resuspended in 0.1% (w/v) Evans blue in TAP. After incubation at room temperature for 5 min, the percentage of stained cells was determined by brightfield microscopy (Axiophot, Carl Zeiss, Germany). At least 500 cells per technical replicate were examined.

**Survival assay**

To obtain the amount of colony forming units (CFUs) of *C. reinhardtii* under the presence of protegencin as a means of algal survival rate, the algal cell density was adjusted to $2 \times 10^6$ cells $mL^{-1}$ in TAP. Subsequently, cells were treated with 0.5% (v/v) DMSO and 0.5% (v/v) protegencin, respectively, for either 1 h or 4 h. An untreated control was handled identically. For the duration of the treatment, the cells were kept shaking in the dark at room temperature. Each treatment was serially diluted in TAP to a final dilution of 1:10,000. 100 µl of each dilution were plated on TAP agar plates in triplicates and incubated at 23 °C for 7 days under a 12:12 LD regime. The percentage of surviving algae was calculated based on the CFUs visible after 7 days.

#### Marker band regions

Raman intensity maps were calculated by summing over compound specific marker bands. We distinguished four components in the analysis namely starch, typical cell compounds, carotenoid and protegencin. Particularly for starch, the analysis included only sharp prominent bands not overlaid by other Raman band features. The three marker bands for typical cell compounds were associated with lipids and proteins with rather broad Raman bands centered at 1451 $cm^{-1}$, 1660 $cm^{-1}$ and 2920 $cm^{-1}$.

The used Raman bands and their corresponding integration regions are listed in Table S1.

#### Segmentation and background correction

Before further analysis of the hyperspectral Raman images, segmentation of the latter in cell and background areas was performed. Furthermore, the previously performed background correction by the SNIP algorithm was supplemented by an additional correction taking the embedding (0.5% agarose) into account. For this purpose, the compound maps of starch, cell, and carotenoid marker bands were normalized on their corresponding maxima and summed up to an intensity map $I_{Cell}(x, y)$ which includes all cell components (Supplementary Fig. 2a). Afterwards the mean value of the obtained intensity map was computed. The mean value served as threshold and pixel values in $I_{Cell}(x, y)$ below or equal to the mean value were assigned as background pixels whereas pixels in $I_{Cell}(x, y)$ with values greater than the mean where assigned cell pixels. Contiguous pixel areas smaller than 100 pixels in the resulting binary cell/background map were assigned to background pixels. The inset in Supplementary Fig. 2b shows an exemplary binary image of the scanned area segmented in algae cell area (red) and background area (blue).

Using the background pixels found, a background correction of the Raman pixel spectra associated with the algal cells was performed to minimize the influence of the background signal originating from the 0.5% agarose embedding. For this, all Raman pixel spectra associated with the background were averaged (see blue spectrum in Supplementary Fig. 2b). This background spectrum was then subtracted from all pixel spectra. Moreover, all single pixel spectra associated with background pixels were then substituted by the newly corrected average background spectrum. Latter was performed to avoid unnecessary fragmentation of the background area by the k-means-cluster-analysis.

The background corrected hyperspectral Raman image served further as the basis for the calculation of the presented Raman intensity maps and were employed for further analysis including k-means cluster analysis, carotenoid detection and the evaluation of the protegencin content in single algae cells.

**K-means cluster analysis**

A k-means cluster analysis with k = 7 was carried out after the above described background correction (3). The spatial cluster arrangement was visualized in false-color-plots after color-coding each cluster.

**Carotenoid detection**

To identify carotenoid clusters in single algae cells a fit of a Lorentzian profile $L(\nu;\nu_0,A,\Gamma) = A\cdot\Gamma/\,[(\nu-\nu_0)^2+\Gamma^2]$ was applied in each of the three marker band regions associated with carotenoid: 1004±20 $cm^{-1}$, 1157±20 $cm^{-1}$, and 1523±20 $cm^{-1}$. Only pixel spectra where all three fits converged were considered for further analysis. We applied an additional constraint to the resulting fit amplitudes $A$: the amplitudes of the Lorentzian profile should be greater than the average spectral intensity plus three times the intensity standard deviation in the wavenumber range 10 $cm^{-1}$ above the corresponding carotenoid spectral region. For instance, the amplitude $A$ computed by a Lorentzian fit model in the wavenumber region of 1004±20 $cm^{-1}$ must be greater than the average spectral intensity plus the three times intensity standard deviation ranging from 1025 $cm^{-1}$ to 1035 $cm^{-1}$. Latter constraint was applied to avoid that pixel with spectral noise are assigned falsely as carotenoid pixels. Pixel spectra fits which did not fulfill these requirements were discarded as possible carotenoid pixels. Contiguous pixel areas smaller than 4 pixels in the resulting binary map were discarded as well.

**Evaluation of the protegencin content**

The metric "Intensity(2160)/Intensity(1451)", defined as the ratio of the integrated spectral intensities of the peak features at 2160 $cm^{-1}$ and 1451 $cm^{-1}$ (see Table S1), served as a measure of the protegencin content. For this purpose, the integrated spectral intensities of the assigned algal cell pixels of the hyperspectral Raman image were averaged and the ratio of the latter was calculated for each investigated algal cell.

**Density Functional Theory calculations of protegencin and its Raman spectrum**

To confirm the detection of protegencin inside *via* Raman Spectroscopy, Density Functional Theory (DFT) calculations were performed on the *trans*-structure of protegencin. The calculations were performed using Orca (version 4.2) (4, 5). First, a relaxed geometry optimization was performed using the hybrid gradient corrected functional B3LYP (6, 7), and polarized triple-ζ basis-set def2-TVZP combined with empirical dispersion correction using the Becke-Johnson damping scheme (8-10). To speed up the calculation resolution of identity approximation for the coloumb integrals with the auxiliary basis set def2/J was used (11, 12).

To confirm that the calculated spectrum represents a minimum in the 3N-6 dimensional potential energy hyperplane and to calculate the Raman spectrum, a subsequent numerical frequency and polarizability calculation was performed using the same functional and basis set, which allows for the calculation of Raman activities.

The calculated Raman activities were converted to Raman intensities by employing the following equation (13):

$$R_i = \frac{(2\pi)^4}{45} \cdot (\nu_0 - \nu_i)^4 \cdot \frac{h}{8\pi^2 c\nu_i \left[1 - e^{-\frac{hc\nu_i}{kT}}\right]} \cdot S_i (1)$$

Herein $R_i, S_i, \nu_i, \nu_0, h, c, T, k$ represent the Raman intensity, Raman activity, vibrational frequency of the ith band, frequency of the incident laser light (in this experiment $3.819 \cdot 10^{14}$ Hz or 785 nm), Planck's constant, the speed of light in vacuum, the temperature (in this case 298K / 25 °C), and Boltzmann's constant. To account for the missing description of anharmonicity and electron correlation in the calculated frequencies, an empirical correction factor of 0.965 was employed (14). To visualize the spectra the calculated line spectrum was broadened using a Lorentzian function with a full-width-at-half-maximum of 18 $cm^{-1}$.

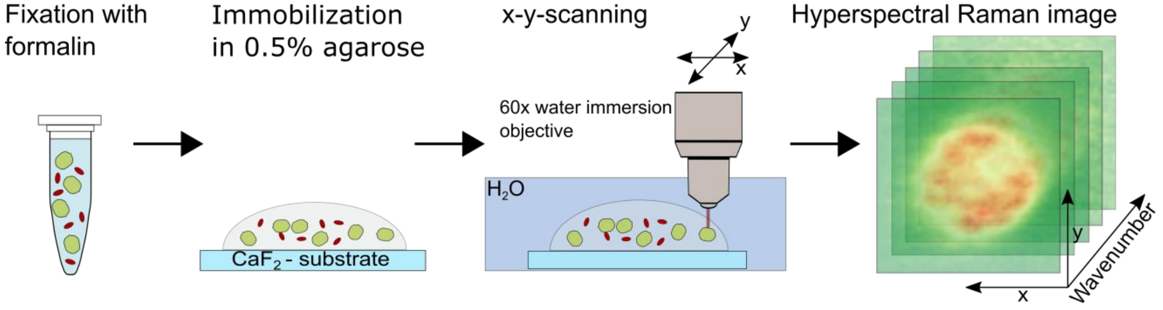


**Supplementary Fig. 1: Schematic of the sample preparation for Raman microspectroscopy and measurement principle.** After the fixation of the algal cells in 4% v/v formalin, the algal cells were immobilized in 0.5% agarose in TAP medium and transferred on a $CaF_2$-platelet. Raman measurements were conducted in aqueous environment and a hyperspectral Raman image was generated by point-wise integration of a 15 µm × 15 µm large scanning grid.

a
b
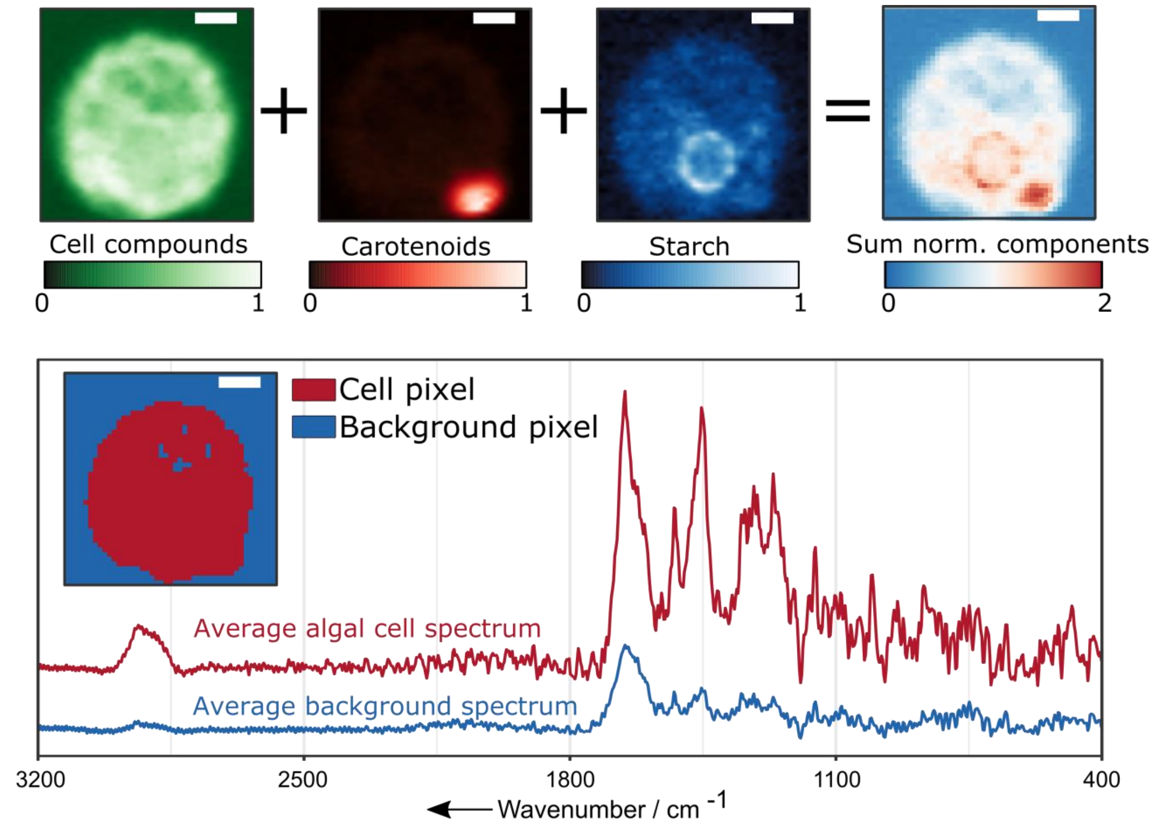


**Supplementary Fig. 2: Raman microspectroscopic image segmentation and calculation of background spectrum. a,** Schematic representation of the workflow to compute an overall intensity map for all cell components by summing the normalized Raman intensity maps of the overall cell compounds, carotenoids, and starch. **b,** Average Raman algal cell (red) and background spectrum (blue) derived from the spectra corresponding to cell and background pixels (see inset for segmentation). Scale bars: 3 µm. For details see Supplementary Information, section segmentation and background correction.

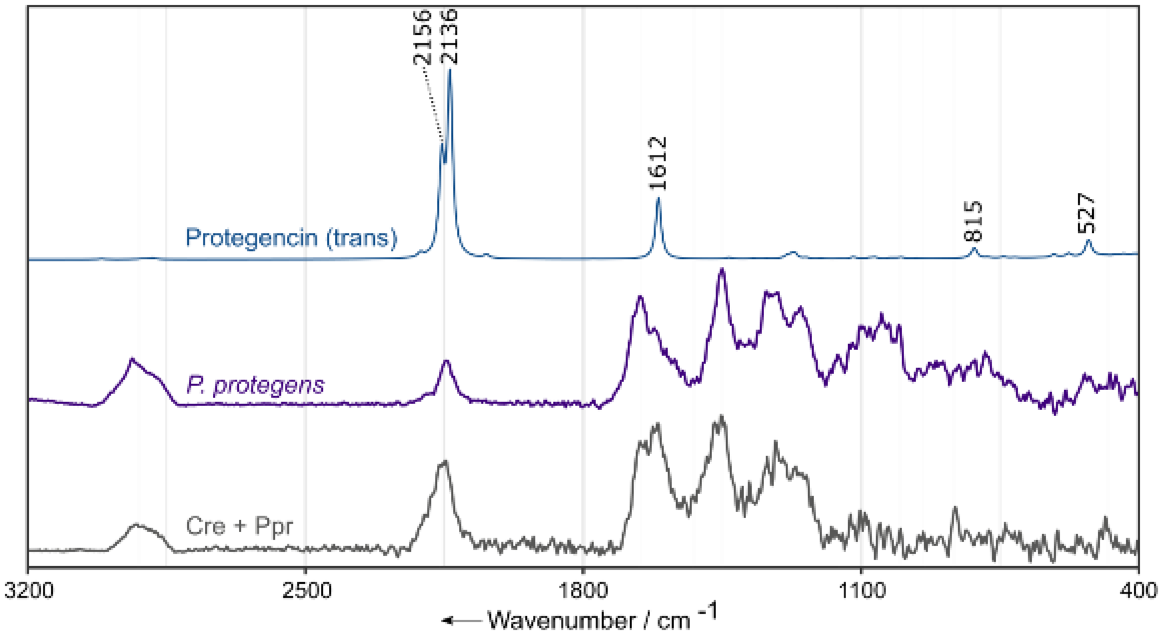


**Supplementary Fig. 3: Comparison of experimental and simulated Raman spectra.** The grey spectrum depicts the average Raman spectrum of an exemplary *C. reinhardtii* cell co-cultivated with *P. protegens* after 16 h incubation time (Cre + Ppr). The purple graph depicts the Raman spectrum of axenic *P. protegens* bacteria after 24 h growth in TAP medium. The theoretical Raman spectrum shown in blue depicts the calculated Raman spectrum of protegencin in *trans*-form.

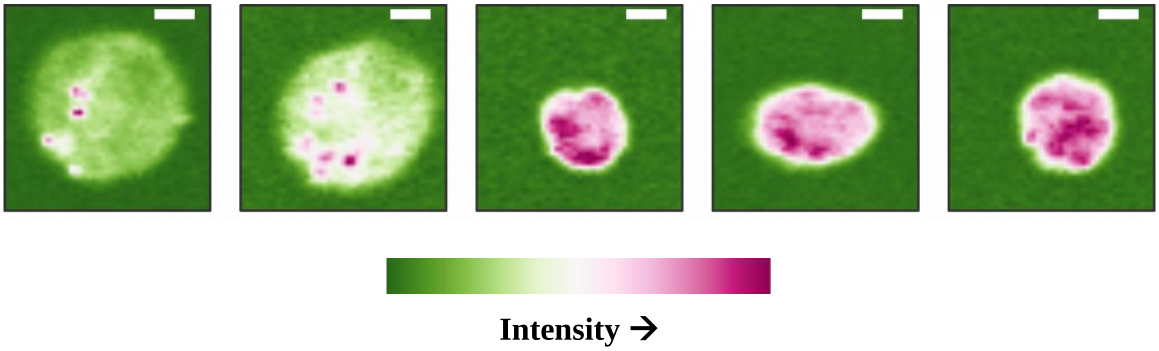


**Supplementary Fig. 4: Distribution of the triple bond substance in five exemplary cells of *C. reinhardtii* after co-culture with *P. protegens* for at least 16 h.** Note: The shown Raman intensities depict the integrated intensities solely after the SNIP background correction to demonstrate that the embedding 0.5% agarose gel gives no contribution to the triple bond peak feature. Scale bars: 3 µm.

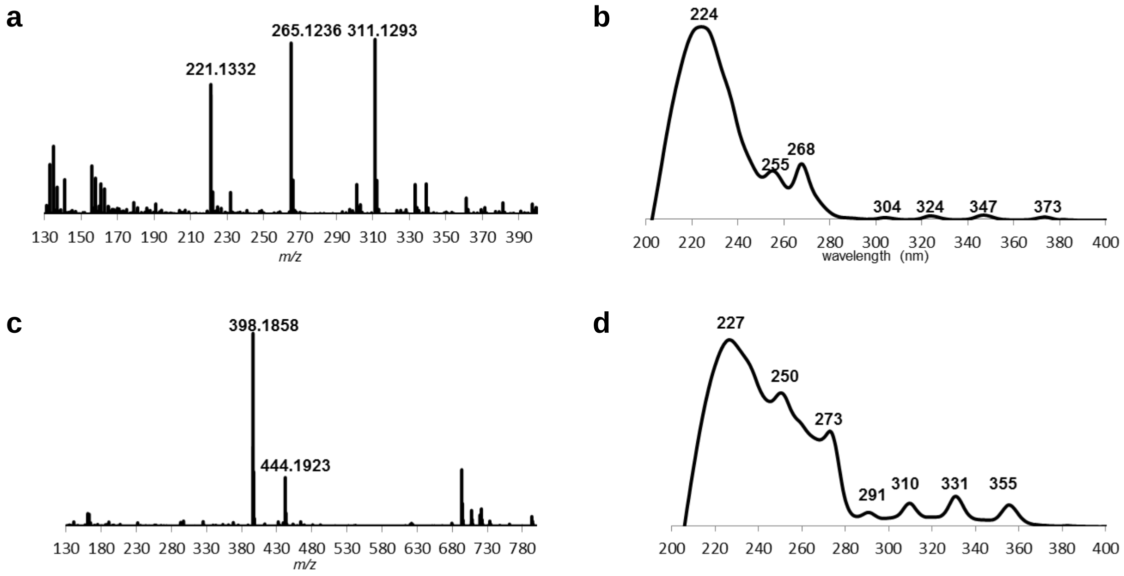


**Supplementary Fig. 5**: **HRMS and UV spectra of protegencin (a and b) ($C_{18}H_{18}O_2$ [M-H]$^-$ calc. 265.1234, obs. 265.1236) and product of its click reaction with benzyl azide (c and d) ($C_{25}H_{25}N_3O_2$ [M-H]$^-$ calc. 398.1874, obs. 398.1858).** Experiments were repeated at least three times.

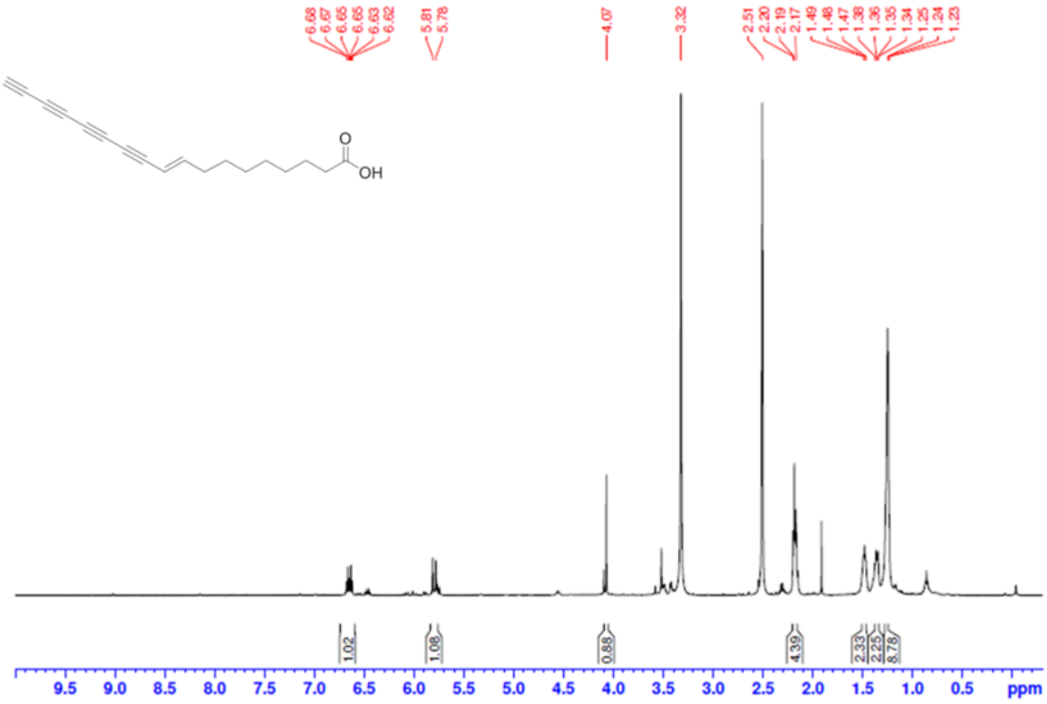


**Supplementary Fig. 6**: **$^1$H NMR spectrum of protegencin.** Experiments were repeated at least three times.

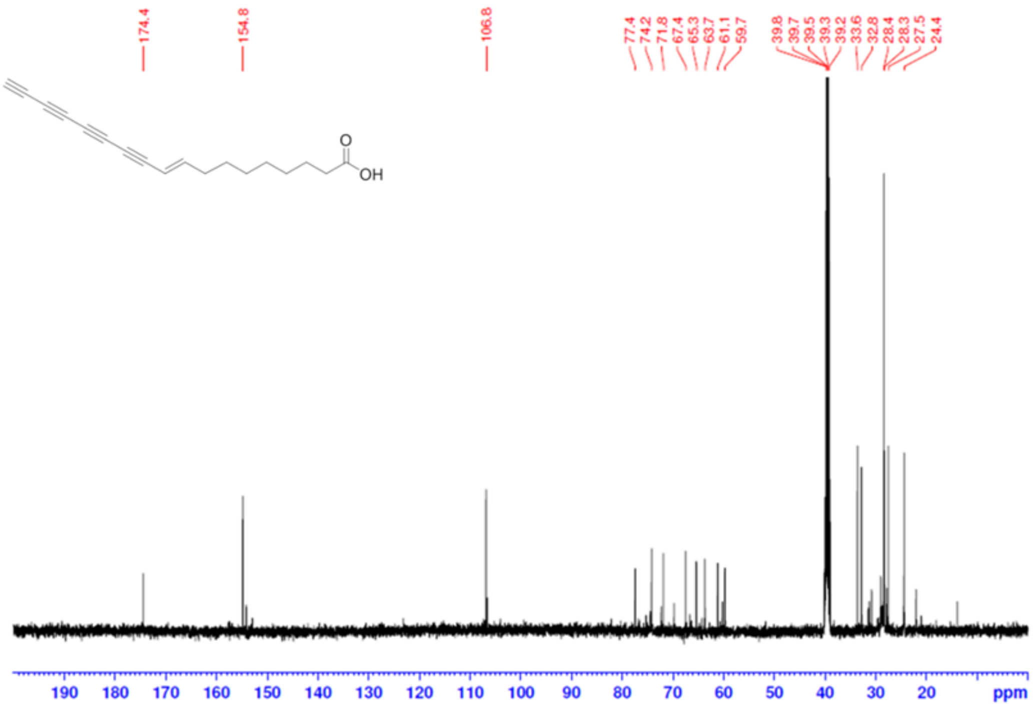


**Supplementary Fig. 7:** $^{13}$**C NMR spectrum of protegencin.** Experiments were repeated at least three times.

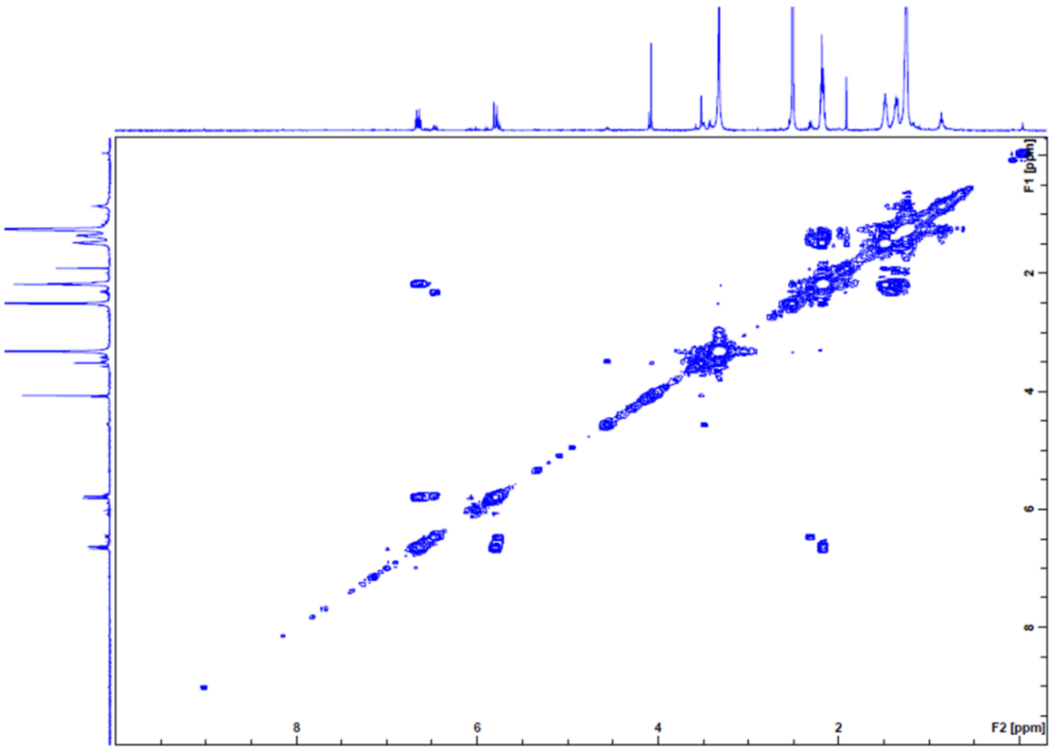


**Supplementary Fig. 8*:* H,H-COSY NMR spectrum of protegencin.** Experiments were repeated at least three times.

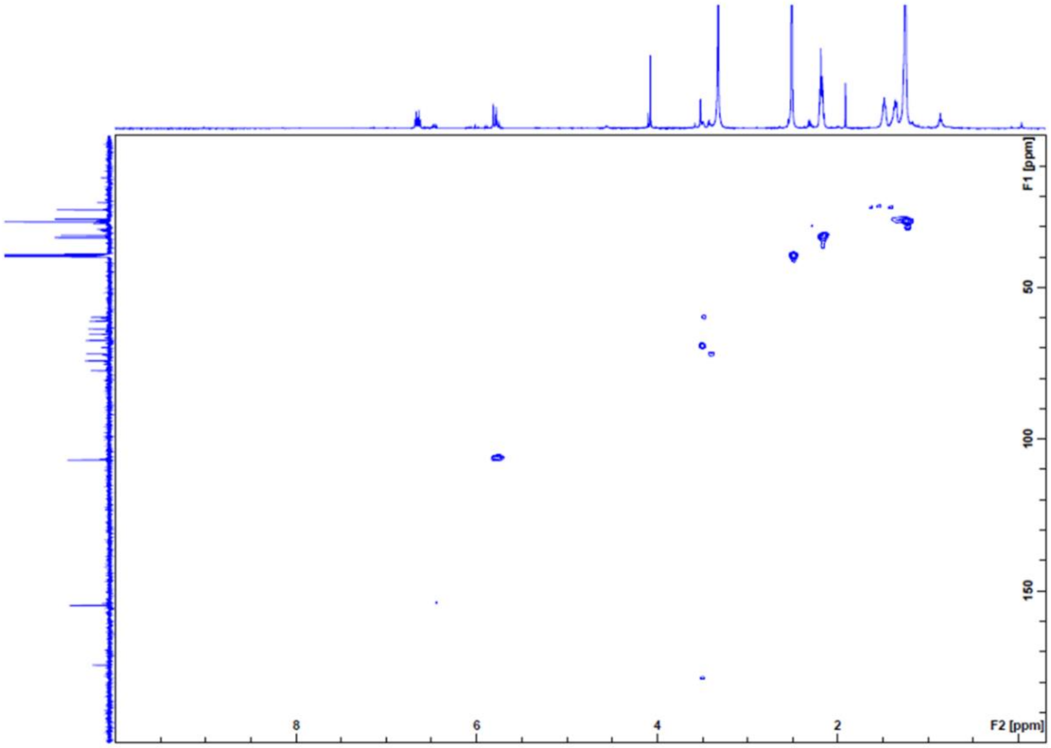


**Supplementary Fig. 9*:* HSQC NMR spectrum of protegencin.** Experiments were repeated at least three times.

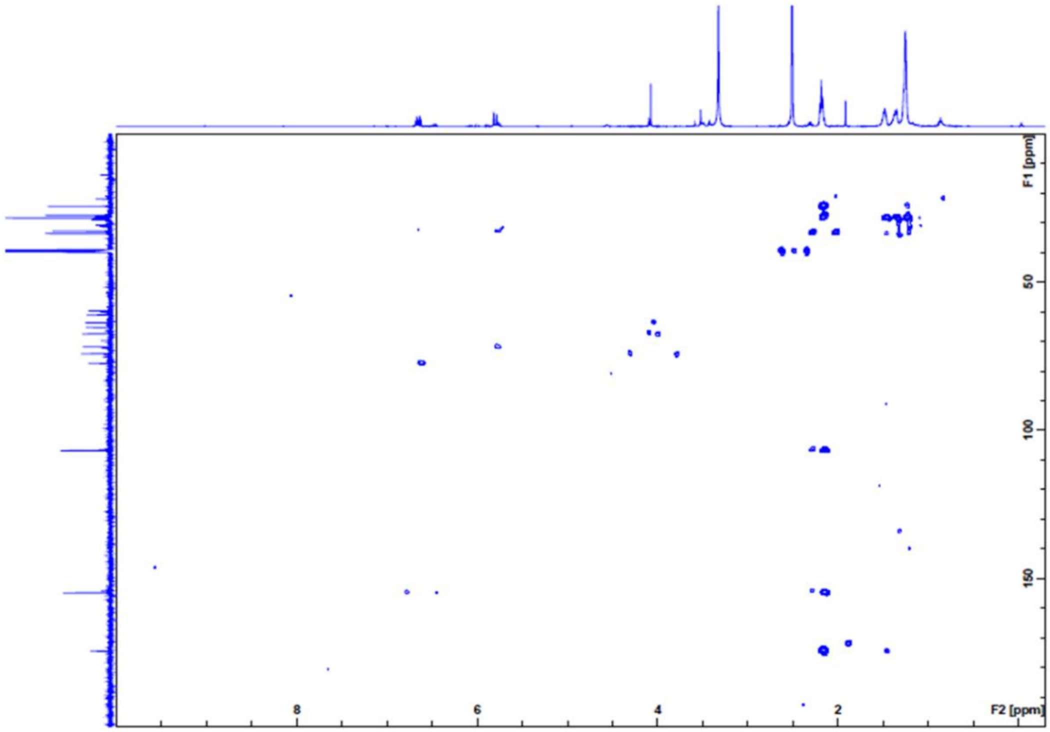


**Supplementary Fig. 10: HMBC NMR spectrum of protegencin.** Experiments were repeated at least three times.

**Supplementary Table 1.** Assignment of the Raman bands of the components of interest with their corresponding integration regions.

| Wavenumber / $cm^{-1}$ | Integration range / $cm^{-1}$ | Assignment | Reference |
|---|---|---|---|
| 478 | ±20 | **Starch**; C–C–C deformation and C–O stretching | (15) |
| 867 | ±20 | **Starch**; C–C–H and C–O–C deformations | (15) |
| 940 | ±20 | **Starch**; C–O–C and C–O–H deformations, C–O stretching | (15) |
| 1004 | ±20 | **Carotenoid**; C–$CH_3$ rocking | (16) |
| 1157 | ±20 | **Carotenoid**; C–C stretching | (16) |
| 1451 | ±30 | **Cell**; Lipids/proteins: $CH_2$ and $CH_3$ deformation | (17, 18) |
| 1523 | ±20 | **Carotenoid**; C=C stretching | (16) |
| 1660 | ±50 | **Cell**; Amide I: C=O stretching, Lipids: C=C stretch vibration | (17, 18) |
| 2160 | ±50 | C≡C stretching | (19) |
| 2920 | ±50 | **Cell**; Lipids/proteins: $CH_3$ symmetric and antisymmetric stretch, $CH_2$ antisymmetric stretching | (17) |

**Supplementary Table 2.** $^{1}H$ and $^{13}C$ NMR data for protegencin (in DMSO-$d_6$).

| **No.** | **$\delta_C$, type** | **$\delta_H$, multiplet, *J* (in Hz)** |
|---|---|---|
| 1 | 174.4, COOH | 11.95, br |
| 2 | 33.6, $CH_2$ | 2.19, t (7.4) |
| 3 | 24.4, $CH_2$ | 1.48, m |
| 4 | 28.4, $CH_2$ | 1.25, m |
| 5 | 28.4, $CH_2$ | 1.25, m |
| 6 | 28.4, $CH_2$ | 1.25, m |
| 7 | 27.5, $CH_2$ | 1.36, m |
| 8 | 32.8, $CH_2$ | 2.19, m |
| 9 | 154.8, CH | 6.65, dt (16.0, 7.1) |
| 10 | 106.8, CH | 5.79, d (15.0) |
| 11 | 77.4, C | |
| 12 | 71.8, C | |
| 13 | 59.7–67.4, C | |
| 14 | 59.7–67.4, C | |
| 15 | 59.7–67.4, C | |
| 16 | 59.7–67.4, C | |
| 17 | 63.7, C | |
| 18 | 74.2, CH | 4.07, s |

**Supplementary Table 3.** Primers for mutation strains of *P. protegens* Pf-5.

| Gene | Forward primer | Reverse primer |
|---|---|---|
| *pgnE Fl1* | 5'- TGGCTGACGGTCGCAACTCCGG -3' | 5'- GCTACTTAATTAAGCTA<u>GCGTAGC</u>CCCTGGATATTGCCGATAAA -3' |
| *pgnE Fl2* | 5'- GCTACGCTAGC<u>TTAATTAA</u>GTAGCATAACCCGCAGCTGGGAG-3' | 5'- GTCGGGCCGGGCCGTTTGACAT -3' |
| $Apr^R$ | 5'-GCTAC<u>GCTAGC</u>ATTCCGGGGATCCGTCGACC-3' | 5'- GCTAC<u>TTAATTAA</u>TGTAGGCTGGAGCTGCTTC-3' |
